# The common origin of symmetry and structure in genetic sequences

**DOI:** 10.1101/199323

**Authors:** G. Cristadoro, M. Degli Esposti, E.G. Altmann

## Abstract

When exploring statistical properties of genetic sequences two main features stand out: the existence of non-random structures at various scales (e.g., long-range correlations) and the presence of symmetries (e.g., Chargaff parity rules). In the last decades, numerous studies investigated the origin and significance of each of these features separately. Here we show that both symmetry and structure have to be considered as the outcome of the same biological processes, whose cumulative effect can be quantitatively measured on extant genomes. We present a novel analysis (based on a minimal model) that not only explains and reproduces previous observations but also predicts the existence of a nested hierarchy of symmetries emerging at different structural scales. Our genome-wide analysis of H. Sapiens confirms the theoretical predictions.

## 1 Introduction

Compositional inhomogeneity at different scales has been observed in DNA sequences since the early discoveries of long-range spatial correlations, pointing to a complex organisation of genome structure [1, 2, 3]. While the origin of these observations has been intensively debated [4, 5, 6, 7, 8, 9, several investigations indicate the ubiquitous patchiness and mosaic-type domains of DNA as playing a key role in the existence of large-scale structures [7, 10, 11]. Another robust statistical observation on genetic sequences reveals the presence of a strand symmetry known as the “Second Chargaff Parity Rule” [12, 13, 14]. Such a symmetry appears universally over almost all extant genomes [15, 16]. In its simplest form, it states that on a single strand the frequency of a nucleotide is approximatively equal to the frequency of its complement [17, 18, 19, 20, 21]. This original formulation has been later extended to the frequency of short (*n* ≃ 10) oligonucleotides and their reverse-complement [21, 23, 24]. While the first Chargaff parity rule [22] (valid in the double strand) was instrumental for the double-helix structure discovery, of which it is now a trivial consequence, the second Chargaff parity rule remains of mysterious origin and of uncertain functional role. Different mechanisms that attempt to explain its origin have been proposed during the last decades [19, 25, 26, 27, 28]. Among them, an elegant explanation [27] proposes that strand symmetry arises as an asymptotic product of the accumulated effect during evolution of inversions and inverted transposition.

Structure and symmetry are in essence two independent observations: Chargaff symmetry concerns the frequency of short oligo-nucleotides (*n* ≃ 10) and thus does not rely on their actual positions in the DNA, while correlations depend on their ordering and are reported to be statistically significant even at large distances (thousands of bases). Therefore, the mechanism that is shaping the complex organization of genome sequences could be, in principle, different and independent from the mechanism enforcing symmetry. However, the proposal of transposable elements [35] as being key biological processes for both statistical properties suggests that they could be the vector of a deeper connection.

In this paper we show that this connection not only explains previous observations, it also predicts new symmetry- structure relationships. Our results, summarised in Fig. 1, connect symmetry and structure at all scales *f* of the DNA using a minimal domain model that accounts for the accumulated effect of the repetitive action of transposable elements. The key ingredient is the reverse-complement symmetry for domain types, a property we show to be a consequence of the action of transposable elements indiscriminately on both DNA strands. Our model accounts for structures (e.g., the patchiness and longrange correlations in DNA) in a similar way as other domain models do, the novelty is that we show the consequences of the biological origin of domains to the symmetries of the full DNA sequence. Our first main finding is that the Chargaff parity rule extends beyond the frequencies of short oligonucleotides. As a consequence it is valid for a larger class of observables (including all those used to characterise the statistical properties of whole genetic sequences) and at much larger scales (including scales where non-trivial structure is present). Several previous unexplained observations of symmetries can thus be understood as different manifestations of our extended symmetry. Our second main finding is that the Chargaff parity rule is not the only symmetry present in genetic sequences as a whole: there exists a hierarchy of symmetries nested at different structural scales. The theoretical predictions of our model are confirmed on the set of chromosomes of Homo Sapiens.

**Figure 1:**
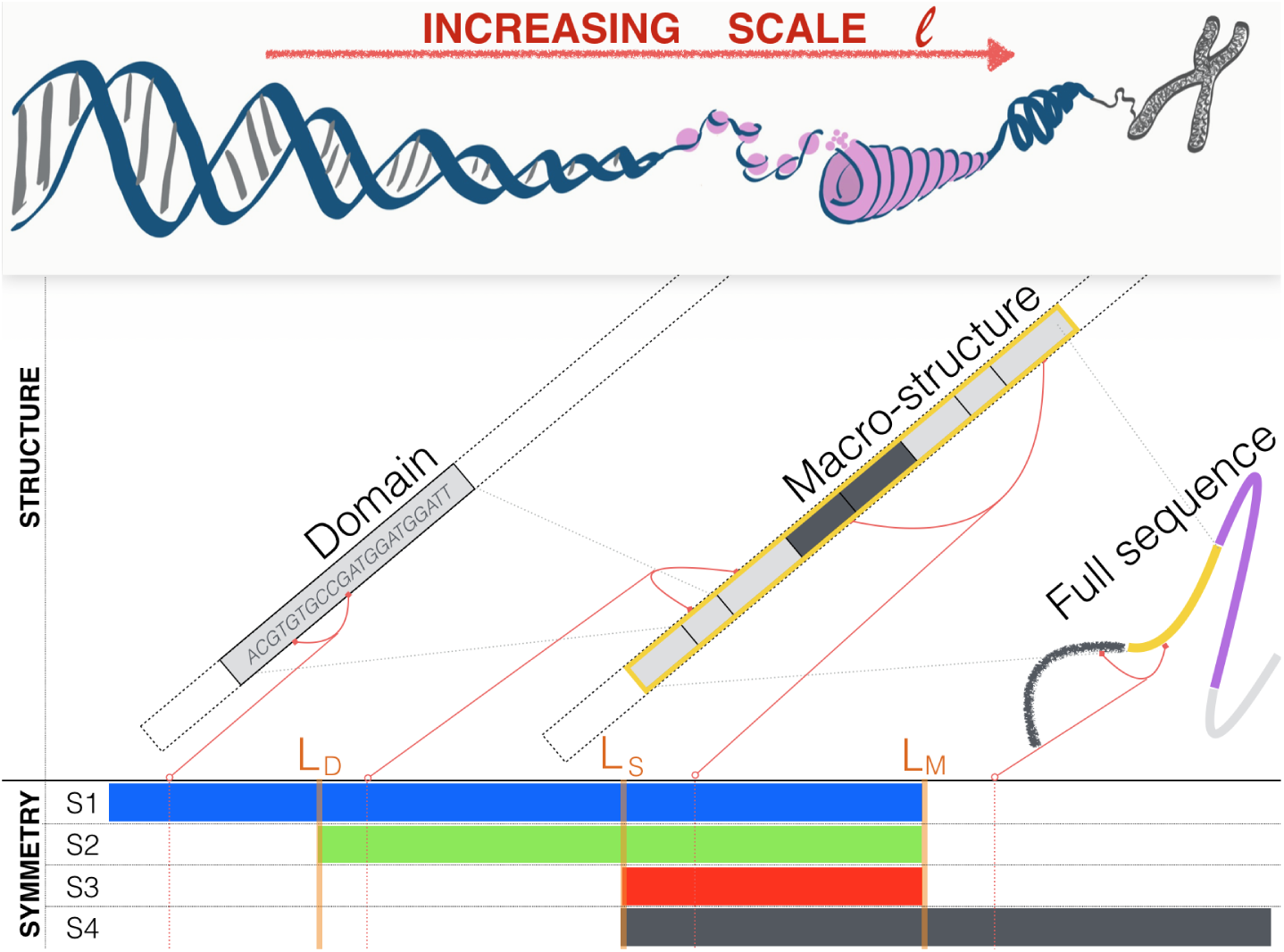
Structure and symmetry at different scales 𝓁 of genetic sequences can be explained using a simple domain model. Our model considers that the full sequence is composed of macro-structures (of size *L*_*M*_) made by the concatenation of domains (of average size *L*_*D*_ *< L*_*M*_), which are themselves correlated with neighbouring domains (up to a scale *L*_*D*_ *< L*_*S*_ *< L*_*M*_). The biological processes that shapes domains imposes that, in each macrostructure, the types of domains comes in symmetric pairs. As a consequence, we show that four different symmetries *S*_1_ *-S*_4_ are relevant at different scales 𝓁 (see text for details).

## 2 Description of the minimal model

We construct a minimal model for DNA sequences s = *α*_1_*α*_2_ *…α_N_*, with *α_i_ ∈*{*A, C, T, G*}, that aims to incorporate the role of transposable elements in shaping statistical features up to the scale of a full chromosome. The model contains three key ingredients at different length scales (see Fig.1 for an illustration):

1. at small scale *L*_*D*_, a **domain** is a genetic sequence **d** = *α*_1_*α*_2_ *α*_*n*_. A domain of a given type is constructed as a realizations (of average size ⟨*n* ⟩ ∼ *L*_*D*_) of a given process *p*. We do not impose a priori restrictions or symmetries on this process. For a given domain type, the symmetrical related type is defined by the process 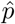 as follows: take a realization (*α*_1_*α*_2_ *… α_n_*) of the process *p*, revert its order (*α*_*n*_α_*n*-1_ *… α*_1_), and complement each base 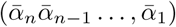 where 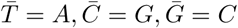
2. at intermediate scale *L_M_* (≫*L*_*D*_), a **macro-structure m** is composed as a concatenation of *m* domains **m** = **d**_1_**d**_2_*…* **d**_*m*_ each belonging to one in a few types. The biological mechanism that motivates our model (transposable elements) imposes that symmetrical related domains (generated by *p* and 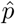) appear with the same relative abundance and size-distribution in a given macro-structure (see SI 5.1 for a justification). We do not exclude that domains of the same type could correlate and thus tend to cluster. We denote *L_S_* the average size of clusters of domains of the same type (*L_D_ < L_S_ ≪ L_M_*);
3. at large scale *L_C_* (*≫ L_M_*) the full sequence is composed by concatenations of macro-structures, each of them governed by different processes and statistics (e.g. different CG content [10] (see also [11]).

## 3 Statistical properties of the model and predictions

We explore statistical properties of typical sequences **s** generated by the domain model described above. We quantify the frequency of appearance in **s** of a given pattern of symbols (an observable *X*) by counting the number of times a given symbol *α*_0_ is separated from another symbol *α*_1_ by a distance *τ*_1_, and this from *α_1_* by a distance *τ_1_*, and so on. More precisely, denote 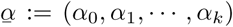 a selected finite sequence of symbols, and by 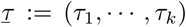 a sequence of gaps. For shortness, we denote this couple by 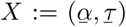 and the *size* of the observable *X* by 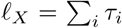 The frequency of occurrence of an observable *X* in the sequence s is

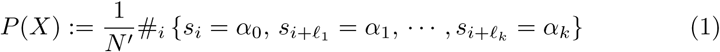

where *N*′ = *N -* 𝓁 _*X*_ + 1 and 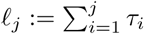. Note that, all major statistical quantities numerically investigated in literature can been expressed in this form, as we will recall momentarily. The main advantage of this formulation is that it allows to inspect both the role of symmetry (varying 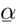) and structure (varying scale separations 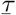) and it thus permits a systematic exploration of their interplay. We say that a sequence has the symmetry *S* at the scale 𝓁 if for any observable *X* with length 𝓁 _*X*_ = 𝓁 we have, in the limit of infinitely long **s**, *P* (*X*) = *P* (*S*(*X*)) where *S*(*X*) is the observable symmetric to *X*. Next, we explore at different scales *f* which symmetries *S* are valid in sequences constructed in agreement with our model.

We start with a natural extension to observables *X* of the reverse-complement symmetry considered by Chargaff. The observable symmetric to 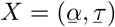 is

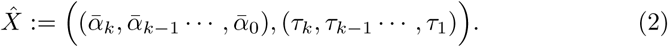

When the size of the observable *X* is smaller than average macro-structure size *L_M_*, the count in Eq. 1 is dominated by occurrences of *X* lying fully inside a macro-structure, as contributions from observation overlapping different macrostructures are negligible. Recalling that in a given macro-structure domain-types appear in symmetric pairs with equal average length and abundance, the symmetry of counts in Eq. 1 follows (see SI for a rigorous derivation). Our first main result follows:

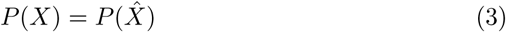

holds for scales 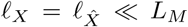 his is an *extension* of Chargaff’s second parity rule because *X* may in principle be an observable involving (a large number of) distant nucleotides and thus Eq (3) symmetrically connects structures even at large scales (up to the scale of macro-structures *L_M_*). By combining *P* (*X*) of different observables *X* we stress that our extended Chargaff symmetry applies to the main statistical analyses investigated in literature, unifying numerous previously unrelated observations of symmetries in the frequency of oligonucleotides^1^, in the autocorrelation function^2^, and in the recurrence-time distribution^3^. These previous results are thus all different manifestations of our extended symmetry, Eq. 3. In view of our model, they can be interpreted as a consequence of an underlying skeleton of symmetric domains shaping the DNA sequence.

We now show that the predictive power of the proposed model goes beyond the clarification and unification of the origin of previous unexplained observations. In order to disentangle the role of different symmetries at different scales 𝓁 we construct a family of observables 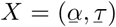 for which we can scan different length scales by varying the gaps vector 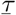. Particularly useful is to fix all gaps in 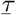 but a chosen one *τ_j_*, and let it vary through the different scales that characterize our model *L_D_, L_S_, L_M_*. To be more specific consider the following construction: given two patterns 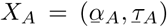 and 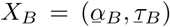 we look for their appearance in a sequence, separated by a distance 𝓁 This is equivalent to look for composite observable 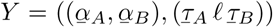or, for simplicity, *Y* =: (*X_A_, X_B_*; 𝓁). We consider two patterns *X_A_, X_B_* of small (fixed) size 𝓁*_X_A,* 𝓁*_X_B ≪L_D_* and we vary their separation 𝓁 to investigate the change in the role of symmetry of domains types and of their compositional organization in the full structure of the sequence: at small 𝓁, *P* (*X_A_, X_B_*; 𝓁) receives the leading contribution from counts fully inside single domains, and thus our extended symmetry Eq. 3 is valid. As 𝓁 grows, the contribution from counts crossing different domains increases. As in our model macro structures are build as a concatenation of different realization of domain-type processes, it predicts that for 𝓁 *≫ L_D_* the same frequency should be expected if the order of *X_A_* and *X_B_* is reversed. Below we investigate the interplay of these two effects at the different scales 𝓁.

The symmetries we consider are defined as compositions of the following two transformations: the first (*R*) reverses the order in the pair by 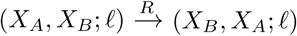 and is relevant as soon as *X_A_* and *X_B_* are in different domains that could equally appear in reversed order; the second (*C*) applies our extended symmetry Eq. 3 to the first of the two observable in the pair by 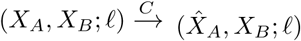 and is relevant when relating counts in symmetrically related domains. Note that *RC* ≠ *CR* (*R* and *C* do not commute), *RR* = *CC* = *Id* (i.e. *R, C* are involutions). *CRC* is the symmetry equivalent to Eq. 2. For a given set *S* of different compositions of *C* and *R*, we denote by (*Y*) the orbit of *Y* under *S* that is the set of different observables obtained acting on *Y* with all combinations of transformations in *S*. For example if *S*1 = {*CRC*} then 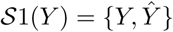 because 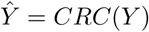 and *Y* = *CRCCRC*(*Y*). We say that a symmetry defined by the set *S* is valid at the scale *l_Y_ ≈ f* if *P* (*Y*) is the same for all observables in *S*(*Y*).

Our model predicts a nested hierarchy of four symmetries *S*_1_-*S*_4_, summarized in Fig. 2, to be valid at different scales 𝓁:

**Figure 2:**
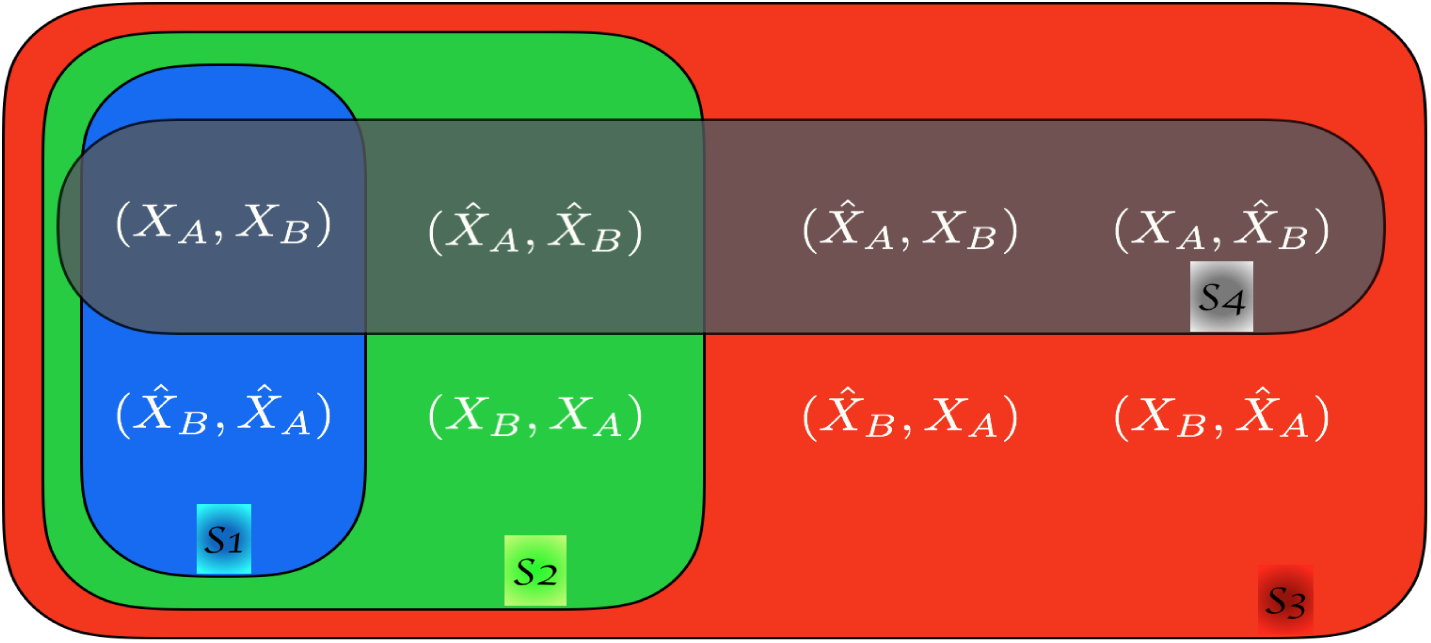
Illustration of the nested sets related by the symmetries *S*1 (blue), *S*2 (green), *S*3 (red), *S*4 (black) for a reference observable *Y* = (*X_A_, X_B_,* 𝓁).

- (𝓁 *≪ L_D_*): *P* (*X_A_, X_B_,* 𝓁) is dominated by *X_A_* and *X_B_* in the same domain. As domain-types appear symmetrically in each macro-structure, 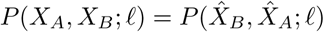 This can be derived directly from Eq.3.

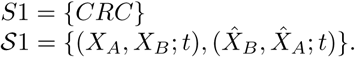
- (*L_D_ ≪ l ≪ L_S_*): *P* (*X_A_, X_B_,* 𝓁) is dominated by *X_A_* and *X_B_* in different domains. As domains are independent realizations, the order of *X_A_* and *X_B_* becomes irrelevant and therefore *R* becomes a relevant symmetry (in addition to *CRC*). If domains of the same type tend to cluster, then for *L_D_ < l ≪ L_S_* the main contribution to *P* (*X_A_, X_B_,* 𝓁) comes from *X_A_* and *X_B_* in different domains of the *same type* (i.e., on different realizations of the same process *p*).

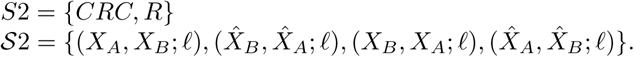 Note that *S*1 *⊂ S*2.
- (*L_S_ ≪* 𝓁 *≪ L_M_*): *P* (*X_A_, X_B_,* 𝓁) is dominated by *X_A_* and *X_B_* in different domains inside the same macro-structure. In addition to the previous symmetries, *C* is valid.

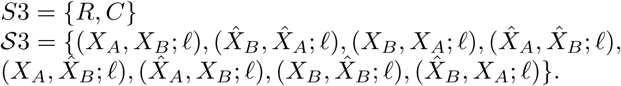 Note that *S*1 *⊂ S*2 *⊂ S*3.
- (𝓁 *≪ L_M_*): *P* (*X_A_, X_B_,* 𝓁) is dominated by *X_A_* and *X_B_* in different macrostructures. Note that the frequency of *X_A_* in one macro-structure and 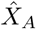 in a different macro-structure are, in general, different. Therefore, for generic *X_A_, X_B_* we have 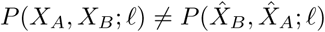 meaning that *S*1 (and thus *S*2 and *S*3) is no longer valid. On the other hand, Eq.3 is valid for both *X_A_* and *X_B_* separately and thus they can be interchanged in the composite observable *Y*. *S*4 = {*RCR, C*}

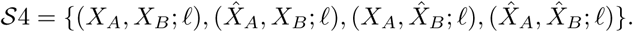 Note that *S*4 *⊂ S*3.

The bottom panel of Fig. 1 illustrates the hierarchy of symmetries at different scales 𝓁 emerging from the analyses above.

## 4 Symmetry and structure in Homo Sapiens

We test our predictions in the human genome. To keep the analysis feasible, we scrutinize the case where *X_A_* and *X_B_* are dinucleotides. This choice has two advantages: by keeping the number of nucleotides in each *X* small we improve their statistics, but at the same time we still differentiate 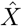 from the simpler complement transformation. Finally, in order to compare results for pairs *X_A_, X_B_* with different abundance, we normalise our observable by the frequencies *P* (*X*) obtaining

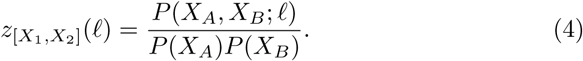

Deviations from *z* = 1 are signatures of structure (correlations). Figure 3 shows the results for chromosome 1 of Homo Sapiens, using a representative case of eight symmetrically related (by *S*1*, S*2*, S*3*, S*4) pairs of dinucleotides. The data shows that at scales 𝓁 *< L_D_* curves appear in pairs (symmetry *S*1) which almost coincide even in the seemingly random fluctuations; around 𝓁 ⋍ *L*_*D*_ two pairs merge forming two groups of four curves each (symmetry *S*2). At larger scales all curve coincide (symmetry *S*3). At 𝓁 *> L_M_* two groups of four observable separates (symmetry *S*4). Similar results are obtained for all choice of dinucleotides and for all chromosomes (see SI: Supplementary data).

**Figure 3:**
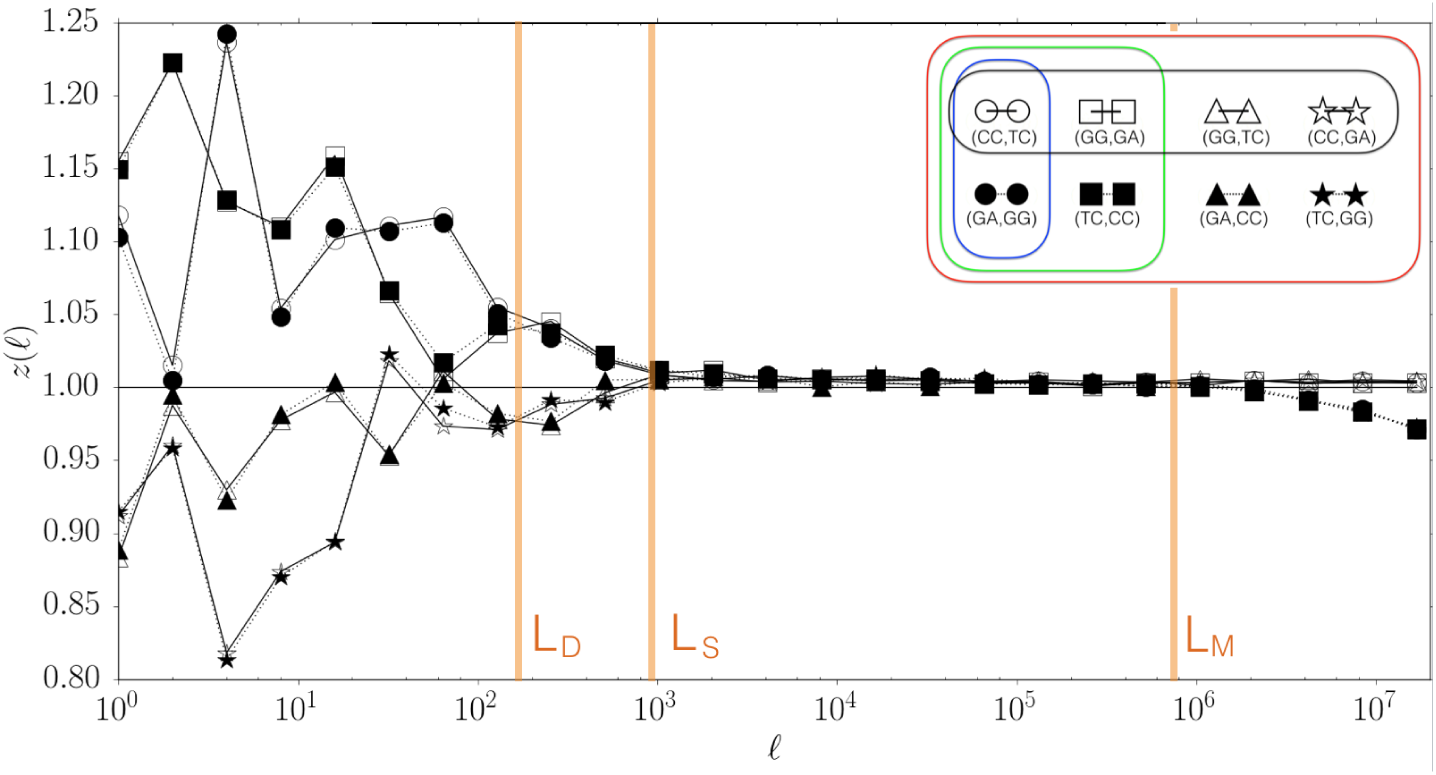
The normalized cross-correlations *z*_[*CC,T* *C*_](𝓁) as a function of the scale 𝓁, together with that of its symmetrical related companions. Note that *S*1*, S*2 and *S*4 are significant even at scales where non trivial correlations are present *z* ≠ 1.

The scale-dependent results discussed above for particular (*X_A_, X_B_,* 𝓁) motivate us to quantify the strength of validity of a symmetry at different scale 𝓁. At this purpose, we compute for each symmetry *S* an indicator *I_S_* (𝓁) that measures the distance between the curves *z*(𝓁) of symmetry-related pairs (*X_A_, X_B_*) and compares it to the ones that are not related by *S* (see Materials and Methods: Quantification of symmetries). Perfect validity of symmetry *S* at scale 𝓁 corresponds to *I_S_* (𝓁) = 0, while *I_S_* (𝓁) = 1 indicates that symmetry-related observables are as different as any two observables. Figure 4 shows the results for chromosome 1 and confirms the existence of a hierarchy of symmetries at different structural scales, as predicted from the analysis of our model (compare color bars of Fig.1 and Fig.4). The estimated relevant scales in chromosome 1 (of total length *N ≈* 2 *×* 10^8^) are *L_D_ ≈* 10^2^, *L_S_ ≈* 10^3^, *L_M_ ≈* 10^6^. As a first independent confirmation of our modelling, note that *L*_*D*_ and *L_M_* are compatible with the known average-size of transposable element and isochore respectively. Moreover, the results for all Homo-Sapiens chromosomes, summarised in Fig.5, show that not only the expected hierarchy is present, but also that the scales *L_D_, L_S_,* and *L_M_* are comparable across chromosomes. This remarkable similarity (see also [36, 37]) is another independent confirmation of our modelling, as it shows that some of the mechanism shaping simultaneously structure and symmetry works similarly in every chromosomes and/or acts across them^4^.

**Figure 4:**
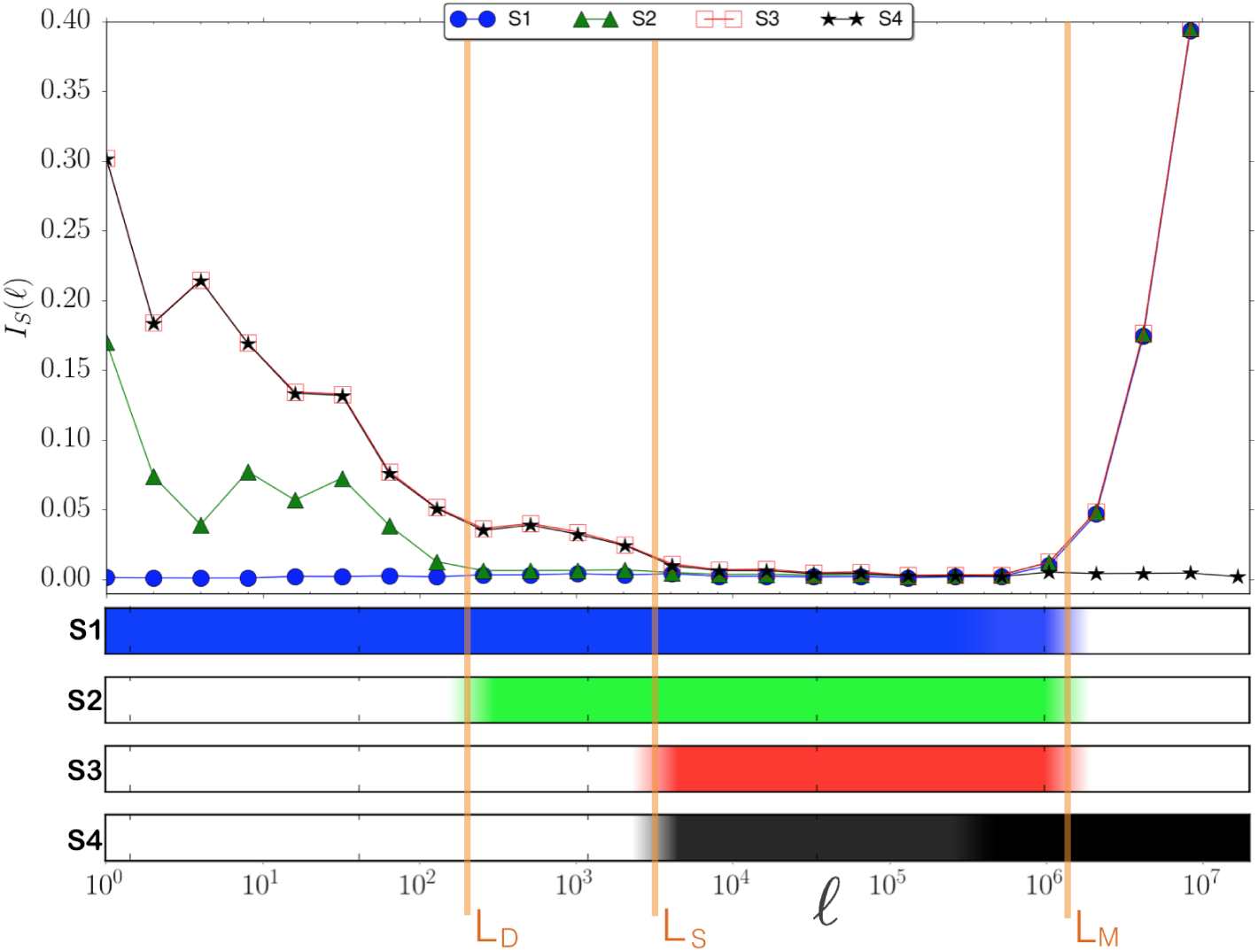
Hierarchy of symmetries in Homo Sapiens. [*Upper panel*] The symmetry index *I_S_* (𝓁) as a function of the scale 𝓁 the smaller the value the larger the importance of the symmetry. [*Bottom panel*] The color bars summarise the onset of the different symmetries: symmetry is considered present if 0 *≤ I_S_ ≤* 0.025 and bar is (linearly interpolated) from full color to white, correspondingly.

**Figure 5:**
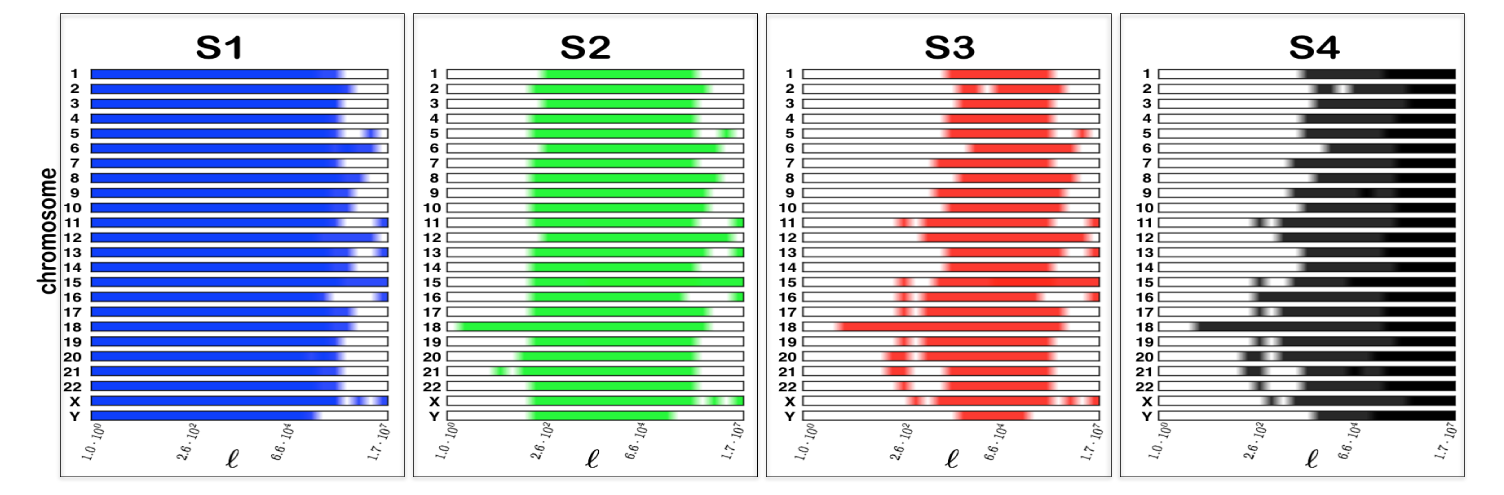
Hierarchy of symmetries in Homo Sapiens. The symmetry index *I_S_* (𝓁) as a function of the scale 𝓁 for the full set of chromosomes in Homo Sapiens. The brighter the color, the larger is the relevance of the symmetry *S*_1_*, S*_2_*, S*_3_, or *S*_4_.

### 4.1 Conclusion and perspectives

At the time of the discovery of the double-helix assembly of DNA, the complement symmetry described by the First Chargaff Parity Rule was used as a key ingredient for the solution of the genetic structure puzzle. What nowadays appears as a trivial consequence of the constitutional property of the molecule, is a demonstration of the fruitfulness of the interplay between symmetry and structure in genetic sequences. In a similar fashion here we show that structural complex organisation of single-strand genetic sequences and their nested hierarchy of symmetry are not features of separate origins; at contrary they should be investigated together as manifestations of the same shaping biological processes. As a consequence, we expect that their concurrent investigation and their interplay will shed a light into their (up to now not completely clarified) evolutionary and functional role. For this aim, we expect that an extension of the analyses presented here to different organisms to be key to clarify and differentiate specific aspects of the mechanism generating the different hierarchy of scales and symmetries at different organism complexity [23]. In parallel, we speculate that the unraveled hierarchy of symmetry at different scale could play a role in understanding how chromatin is spatially organised if investigated in conjunction with the puzzling functional role of long-range correlations [38].

## Materials and Methods

### Quantification of symmetries

For a given reference pair *Y*_ref_ = (*X_A_, X_B_*), and a fixed symmetry *S ∈ S*1*, S*2*, S*3*, S*4 we consider the following distance of *Y*_ref_ to the set *S*

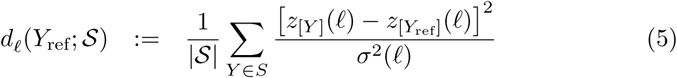

where *σ*(𝓁) denotes the standard deviation of *z*(𝓁) over all *Y*. We then average over the set of *A* all *Y*_ref_ to obtain a measure of the strength of symmetry *S* at scale 𝓁 given by the index

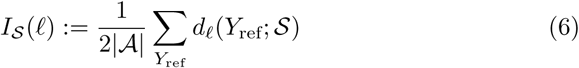

Note that *I_S_* (𝓁) = 0 indicates that *z* is the same for all *Y* in the set *S* and thus indicate full validity of the symmetry *S* at the scale 𝓁. At contrary *I_S_* (𝓁) = 1 indicate that *z* variates in as *S* much as in the full set, at the scale 𝓁, indicating thus that *S* is not valid at that scale.

### Data handling

Genetic sequences of Homo Sapiens were downloaded from the National Center for Biotechnology Information (ftp://ftp.ncbi.nih.gov/genomes/Hsapiens). We used reference assembly build 38.2. The sequences were processed to remove all letters different from *A, C, G, T*.

## Acknowledgements

EGA and GC thank the Max Planck Institute for the Physics of Complex Systems in Dresden (Germany) for hospitality and support at the early stages of this project.

## 5 Supplementary Information

### 5.1 Dynamics leading to symmetric domain model

We show that the cumulative action of a reverse-complement cut-and-paste mechanism creates symmetrically-related domains.

For concreteness, consider the following three steps that encapsulate the combined effect of different shaping forces (i.e. transposable elements) in the evolution of a genetic sequence as a whole:

#### • Initialization

1. Start from a sequence **m**(*t* = 0) of length *N* generated by a process γ such that **m** has frequencies not obeying Second Chargaff Parity Rule, that is *f*γ(*A*) ≠ *f*γ(*T*) and *f*γ(*C*) =≠ *f*γ(*G*).

Note that ┌ is an arbitrary process and it can in principle generate a sequence **m** with a complex structure (e.g., domains and clusters of CG content) or be a simple stochastic process (e.g., a Markov model of low order).

#### • Dynamical evolution

The sequence **m** evolves in time *t* ⟼ *t* + 1 through the repetitive application of two steps:

2. Choose a length *l* from a distribution *ρ*(*l*).

3. Choose at random in **m** a subsequence of length *l* and replace it with its reverse-complement.

We choose *ρ*(*l*) such that it has a well-defined mean 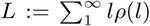 and that 1 *≪ L ≪ LM*.

We now characterise the properties of the sequence **m**(*t*) as we increase the number of iterates *t* of the dynamics described above. We say that the site *αi ∈* **m**(*t*) is of type 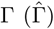 if at time *t* it participated an even (odd) number of times in the reversecomplement event in step 3 above. We investigate the length in **m**(*t*) of subsequences (domains) of consecutive sites of the same type. The average size of such domains after *t* iterations of the dynamics is denoted by ⟨*l*⟩_γ_(*t*) and 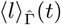 for the two domain types respectively.

Three different regimes in time can be identified:

i. At short times *t < t*_overlap_ = *L_M_ /L* the subsequences involved in step 3 are all distinct and therefore we estimate

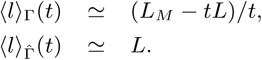 Note that 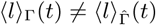 and the sequence is not Chargaff symmetric.
ii. At intermediate times *t*_overlap_ *< t < t*_random_ = *L_M_* the subsequences involved in step 3 overlap. In this regime, the average domain size of the two types are equal and decrease in time as:

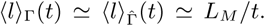 The sequence is structurally complex with domains of average size 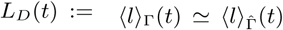 The equal representation of domain types implies that the sequence is Chargaff symmetric.
iii. At large times *t > t*_random_ the dynamics reach equilibrium and

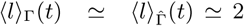 The sequence is random (no structure) and Chargaff symmetric.

The intermediate regime is robust (large average life-time as the ratio *t_random_/t_overlap_* ≃ *L >>* 1) and it is connected to the biologically relevant cases, as complex structure and symmetry coexist (symmetric domain model). Different macro-structures **m** with different statistical properties can be combined at a larger scale^5^.

### Derivation of the nested hierarchy of symmetries

We derive the nested hierarchy of symmetries in the minimal model for genetic sequence. To fix notations, we describe our model and its statistical properties as follows:

- The full sequence is build concatenating *r* macro-structures: **s** = **m**1 **m**2 *…* **m***r*.
- A macrostructure **m** is build concatenating *m* domains: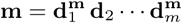
- The average domain length^6^ is denoted by *LD*, the average macro-structure length is denoted by *LM*. The total length of the sequence is *N*
- A domain **d^m^** in the macro-structure **m** is a finite-size realisation of a process chosen between two^7^ symmetrically related process-types: *C***_m_** and 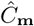 The average size of domains is *LD*. We use the notation **d** *∈ C* to indicate that **d** is generated by the process of type *C*.
- For a a given observable X, we denote by *fC* (*X*) the limiting relative frequency^8^ of occurrence of *X* in a domain of type *C*. Recall that, the definition of symmetrically related processes (of the same macro-structure) imposes that, for every choice of *X*:

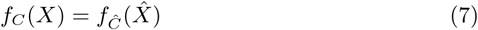 At contrary, different macro-structures have in principle different process-types statistics.
- We denote by 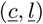 an ordered sequence of domains of types 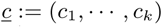; c_j_ϵ 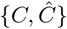 of lengths 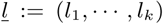 respectively; and by 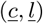 the sequence of domains defined by 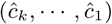 and (*l_k_, …, l*_1_). We denote by 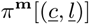 the relative frequency^9^ of counts of subsequence of domains 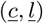 in **m**. We use the shortcut by *π***^m^**({*c, l*}) = *π*[(*c, c, …, c*), (*l*_1_*, l*_2_*, …, l_k_*)] with 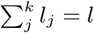 to denote the relative frequency of a cluster of length *l* of domains of the same type *c*. For 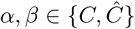 and *k ≥* 1, we denote *π***^m^**(*α, β*; *k*) the relative frequency of *j* such that **d***_j_ ∈ α* and **d***_j_*+*k ∈ β*.
- In each macro-structure, the probability distribution of domain-sizes is denoted *p***m**(*l*).
- We do not enforce any prescription to concatenate domains in a macrostructure (determined by *π*), but the following properties:
  *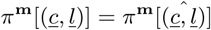* This ensure that the structural statistics of two symmetrically coupled domain-types ordering is unbiased.
  for *k >> LS/LD*; *π***^m^**(*c*1*, c*2*, k*) = *π***^m^**(*c*1)*π***^m^**(*c*2). This defines the average length *LS* beyond which correlations in domain ordering can be neglected. *LS* is thus the average size of clusters of domains of the same type.

We start by showing our first main result: we denote by 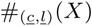 the counts of *X* inside 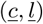 Using 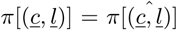 and *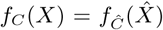*we have that 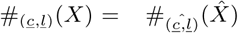 Finally, for *X* of size 𝓁*_X_ ≪ L_M_*, the counts of *X* in the full sequence is dominated by *X* not overlapping different macro-structures and thus we conclude that

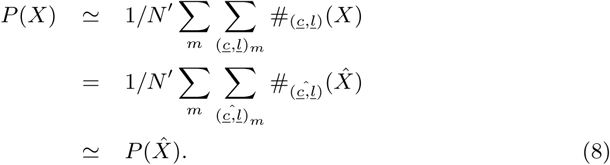

We proceed showing the nested hierarchy of symmetries. In the following, we always approximate the counts of *X* inside a domain of type *C* and of length *l* by *l fC* (*X*) as we consider observable *X*_*A*_ and *X*_*B*_ of size much smaller than typical domain sizes *LD*.

Define

#_(*i*)_(*Y*) := number of *Y* fully inside the *i*-th domain

#_(*ij*)_(*X_A_, X_B_,* 𝓁):= number of *X*_*A*_ fully in the *i*-th and *X*_*B*_ in the *j*-th domains, at distance 𝓁

#(*Y*):=number of *Y* := (*X, XB,* 𝓁) in the full string

=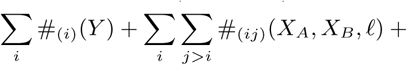+{terms where *X*_*A*_ or *X*_*B*_ overlap domains boundaries} As we will consider only the case *lXA, lXB ≪ LD*, we neglect the last term.

- (𝓁 *≪ LD*): At these scales the following sum dominates,

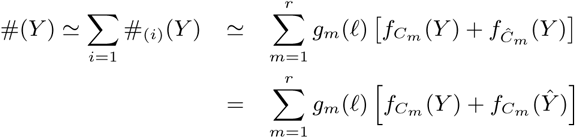

where

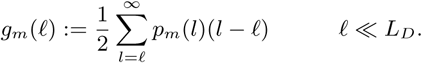

We conclude that 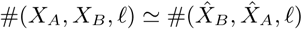 at these scales, and thus symmetry *S*1 is valid. This can also be derived directly from Eq. 8.

For 𝓁 *>> LD*, *X*_*A*_ and *X*_*B*_ typically lie in different domains and therefore the second term in Eq. 9 dominates

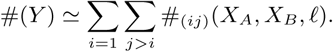

The counts will be estimated as the product of the probabilities of *X*_*A*_ and *X*_*B*_ because each domain is an independent realisations. At different scales *£* there are different relationships between the domains in which *X*_*A*_ and *X*_*B*_ typically lie, leading to the following different cases:

- (*L_D_ <<* 𝓁 *< L_S_*): At these scales the sum is dominated by counts of *Y* inside a cluster of domains of the same type. Each cluster contribute to the counts of *Y* with a term *π*[{*c, l*}](*l -*𝓁)*fC* (*X*_*A*_)*fC* (*X*_*B*_) and thus, in this case

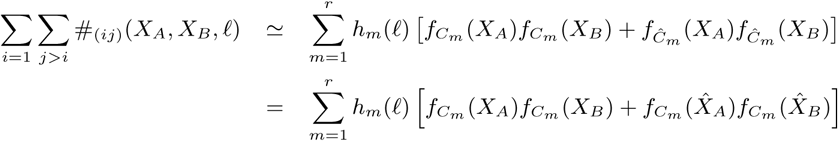

where

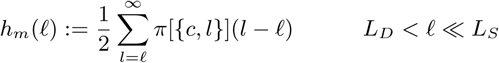

We conclude that 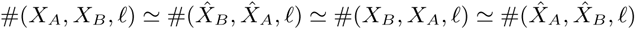 at these scales, and thus symmetry *S*2 (and *S*1) is valid. If the processes are such that correlations inside domains vanishes at a scale smaller than *LD*, then *S*2 sets in at this scale.

- (*L_S_ ≪ -*𝓁 *≪ L_M_*): At these scales the sum is dominated by *X*_*A*_ and *X*_*B*_ lying in different cluster

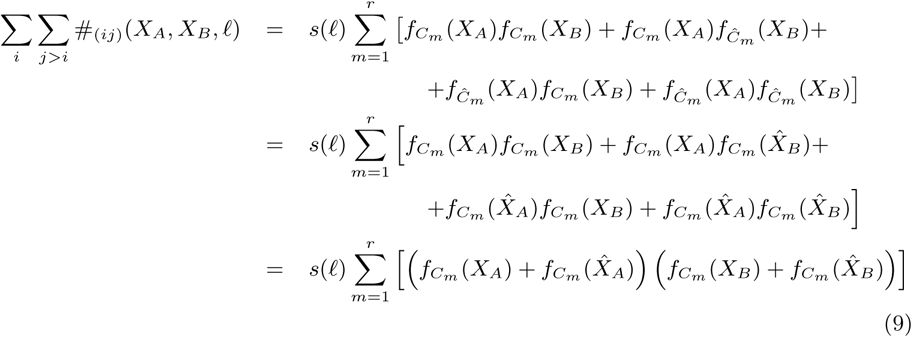

where

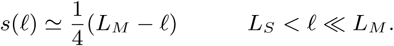 We conclude that 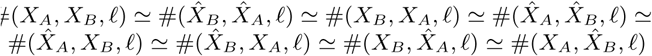 at these scales, and thus symmetry *S*3 (and *S*2*, S*1 and *S*4) is valid.
- (*LM ≪* 𝓁): At these scales the sum is dominated by counts where *X*_*A*_ and *X*_*B*_ are in different macro-structures:

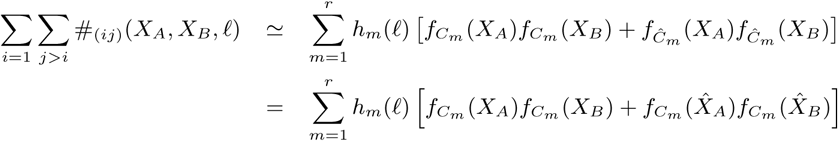

where *qm,n*(𝓁) counts how many sites separated by 𝓁 lie in macro-structures m and n, respectively. We conclude that 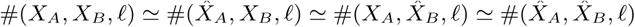 and thus symmetry *S*4 is valid.

### Supplementary data

Data of the values of *z*(*X_A_, X_B_,* 𝓁) for the full set of dinucloetides and for all chromosomes of Homo Sapiens can be visualised and downloaded at the git repository: https://dx.doi.org/10.5281/zenodo.1001805

## Footnotes

1 the frequency of a given oligonucleotide 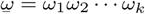 can be computed as *P*_*s*_(*X*) with the choice 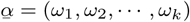 and *τ* = (1, 1, 1) Eq. (3) implies that the frequency of an oligonucleotide is equal to the frequency of its reverse-complement symmetric and thus implies the second Chargaff parity rule, a features that has been extensively confirmed. Note that, the validity of Second Chargaff Parity rule at small scale is not enough to enforce Eq. (3) for generic observable *X* of size up to *L*_*M*_.

2 the autocorrelation function *Cω* (*t*) of nucleotide *ω* at delay *t* is the central quantity in the study of long-range correlations in the DNA. It corresponds to the choice 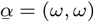 and 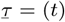. Equation (3) predicts the symmetry 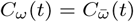 for all *t* ⪡ *L*_*M*_. In the specific case of dinucleotides, such relation has been remarked in Ref. [31]. Our result holds for any oligonucleotide *ω*.

3 the distribution *Rω* (*t*) of first return-time between two consecutive appearances of the oligonucleotide *ω* is studied in Refs. [32, 33]. *Rω* (*t*) can be written as a sum of different *P* (*X*). Eq. 3 predicts 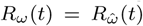 for all *t*⪡ *τ*_*M*_. This symmetry was observed for oligonucleotides in Ref. [34].

4 as an example, it is known that crossing over takes place between homologous transposons located also at different chromosomes.

5 The concatenation of macro-structures can be thought as the result of a dynamic similar to the one described above but acting at larger scales, and with a proper time-scale in the first regime, where no symmetries is enforced between macro-structures.

6 It corresponds to 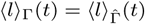 in the previous section.

7 Generalisations to more than two symmetrically domain-types is straightforward and it is not expected to change the main features of the model.

8 We assume that process types *C* are such that *π*_*C*_ (*X*) are well defined for all choice of observables *X* in the limit of size of domains going to infinity.

9 We assume that the structural properties of a given macro-structure is such that *π* is well defined in the limit of number of domains *m* going to infinity.

